# Machine Learning-Enhanced Extraction of Protein Signatures of Renal Cell Carcinoma from Proteomics Data

**DOI:** 10.1101/2025.02.17.638651

**Authors:** Hongyi Liu, Zhuo Ma, T. Mamie Lih, Lijun Chen, Yingwei Hu, Yuefan Wang, Zhenyu Sun, Yuanyu Huang, Yuanwei Xu, Hui Zhang

**Author notes:** Corresponding author. (Hui Zhang). These authors contributed equally to this work (Hongyi Liu, Zhuo Ma).

## Abstract

In this study, we generated label-free data-independent acquisition (DIA)-based liquid chromatography (LC)-mass spectrometry (MS) proteomics data from 261 renal cell carcinomas (RCC) and 195 normal adjacent tissues (NAT). The RCC tumors included 48 non-clear cell renal cell carcinomas (non-ccRCC) and 213 ccRCC. A total of 219,740 peptides and 11,943 protein groups were identified with 9,787 protein groups per sample on average. We adopted a comprehensive approach to select representative samples with different mutation sites, considering histopathological, immune, methylation, and non-negative matrix factorization (NMF)-based subtypes, along with clinical characteristics (gender, grade, and stage) to capture the complexity and diversity of ccRCC tumors. We used machine learning identified 55 protein signatures that distinguish RCC tumors from NATs. Furthermore, 39 protein signatures that differentiate different RCC tumor subtypes were also identified. Our findings offer an extensive perspective of the proteomic landscape in RCC, illuminating specific proteins that serve to distinguish RCC tumors from NATs and among various RCC tumor subtypes.

## Introduction

Renal cell carcinoma (RCC) ranks in the top 10 most frequently diagnosed cancers with an estimated 81,610 diagnoses and 14,390 deaths in the United States in 2024^1,2^. The World Health Organization (WHO) listed 7 RCC subtypes defined by specific molecular aberrations in 2022^3^. Clear cell RCC (ccRCC) is the most predominant subtype and accounts for the majority (75%) of RCC-related deaths^4^. Non-clear cell RCC (non-ccRCC) represents around 25% of RCC, encompassing various rare subtypes predominantly characterized by histopathological properties^3,5,6^. Understanding ccRCC oncogenesis has been greatly aided by The Cancer Genome Atlas (TCGA) project’s comprehensive genomic, epigenomic, and transcriptomic profiling^7,8^. Dysfunctional regulation of the VHL gene and subsequent aberrations related to genes PBRM1, SETD2, KDM5C, or BAP1 are essential for disease advancement and correlated with more aggressive phenotypes^9–11^. Although previous work on non-ccRCC has discovered several genomic changes to aid in differential diagnosis of different RCC subtypes, due to the heterogeneity of non-ccRCC subtypes, genomic features related to non-ccRCC are rarely found^12–15^. Compared with the genomics, proteomics can provide more extensive information corresponding to the occurrence and development of cancer^16–19^. More importantly, protein abundance cannot be reliably predicted from DNA- or RNA-level measurements^20–23^. Therefore, proteomics would be useful for finding common protein signatures between ccRCC, while non-ccRCC distinguishing tumor tissues from normal tissues.

As part of our efforts within the Clinical Proteomic Tumor Analysis Consortium (CPTAC), we have conducted proteomic analyses of RCC using data independent acquisition (DIA)-mass spectrometry (MS). This involved three RCC cohorts^24–26^. DIA is a MS-based proteomics technique that aims to comprehensively and reproducibly record all peptide precursors and their fragments within a given mass range^27–30^. This contrasts with the data-dependent acquisition (DDA), where only the most abundant peptide precursors are selected for fragmentation^31^. While DDA relies on the stochastic nature of peptide precursor selection, which can lead to missing data across multiple runs, DIA instead systematically fragments all precursors within a specified mass range, thereby generating a more comprehensive and reproducible dataset^27,32^. Given the heterogeneous nature of RCC, a technique like DIA is crucial for understanding the molecular basis of the RCC.

In this study, we leveraged the high-throughput, DIA LC-MS to analyze RCC proteome. We performed proteomic profiling of 261 RCC samples and 195 normal adjacent tissues (NAT). The RCC tumors included 48 non-ccRCC and 213 ccRCC. It is worth noting that the 213 ccRCC samples accounts for most of the total RCC samples^24–26^. If all ccRCC and non-ccRCC samples are analyzed together, it will likely obscure or weaken the unique characteristics of non-ccRCC subtypes. Directly extracting all ccRCC samples for comparative analysis failed to reveal the similarities between ccRCC and non-ccRCC. Therefore, selecting representative ccRCC samples for subsequent multi-level analysis can improve the ability to focus on non-ccRCC subtypes while retaining the overall characteristics of RCC. By optimally selecting ccRCC samples representing different mutation sites and pathological types, stages, grades, etc., the diverse characteristics of ccRCC can be displayed to the maximum extent and provide clues for further segmentation. Analysis of the representative ccRCC and non-ccRCC samples can more clearly and systematically reveal their unique properties and functions while retaining complete RCC information.

To achieve this, artificial intelligence approaches such as deep learning and machine learning (ML) methods have demonstrated significant promise in the analysis and interpretation of large-scale proteomic data^33–35^. ML can identify complex patterns in the proteomic data that may be missed by traditional statistical approaches^36,37^. Specifically, ML can assist in identifying protein signatures associated with different RCC subtypes, thereby potentially improving differential diagnosis and contributing to a better understanding of the RCC molecular basis^38,39^.

In this study, we utilized 261 RCC samples and 195 NATs to establish protein signatures that can identify RCC tumor subtypes and distinguish the RCC tumors from NATs by ML. These protein signatures could be used to improve diagnostic accuracy, inform treatment strategies, or even identify potential new therapeutic targets.

## Result

### Proteomic analysis revealed distinct protein expression patterns

We examined proteomics data from a wide range of 48 non-ccRCC tumors and 213 ccRCC tumors and 195 NAT samples. DIA-based proteomic analysis was used to profile all samples for the proteome. The mutation site data were available for 259 of the tumor samples. The associated clinical data and metadata are provided in Table S1 and summarized in Figure S1A.

Since the samples came from different cohorts, to avoid the batch effect, we used block randomization and interspersed NCI-7 quality control (QC) and pool QC samples between the RCC and NAT samples as MS QC to evaluate the robustness of label-free quantification. Tissue type, gender, grade, stage, age, and loading volume were considered during the randomization. A total of 456 samples were divided into 19 sets, each set contained 24 samples. On average, each set contained 14 tumors, 10 NATs, 1 pooled sample QC, and 1 NCI-7 QC. A total of 219,740 peptides and 11,943 protein groups were identified for proteomic study. On average, 9,787 protein groups were detected per sample. Spearman’s correlation coefficients were calculated for the NCI-7 QC samples with an average correlation of 0.99 among the samples. A similar outcome was observed for the pool QC samples. These results demonstrated the consistent stability of the MS platform (Figure S1B).

To visualize proteomic differences across each subtype of RCC tumors, we performed uniform manifold approximation and projection (UMAP) analysis, which visualizes the high-dimensional proteomic data in a reduced-dimensional space and detects patterns and variations in protein expression across RCC tumor subtypes. The resultant UMAP plot displays the RCC subtypes and NATs in different colors (Figure 1A). The results showed a clear separation between RCC tumors and NATs, ccRCC tumors clustered together and separated from NATs, while Oncocytoma type 1, Oncocytoma type 2, and Oncocytoma variant were in one cluster which was far away from ccRCC tumors and NATs. In contrast, the pRCC type 1 and 2 were located between ccRCC tumors and NATs. From the UMAP, the proteomic heterogeneity was clearly indicated between non-ccRCC tumors compared with ccRCC tumors.

**Figure 1.**
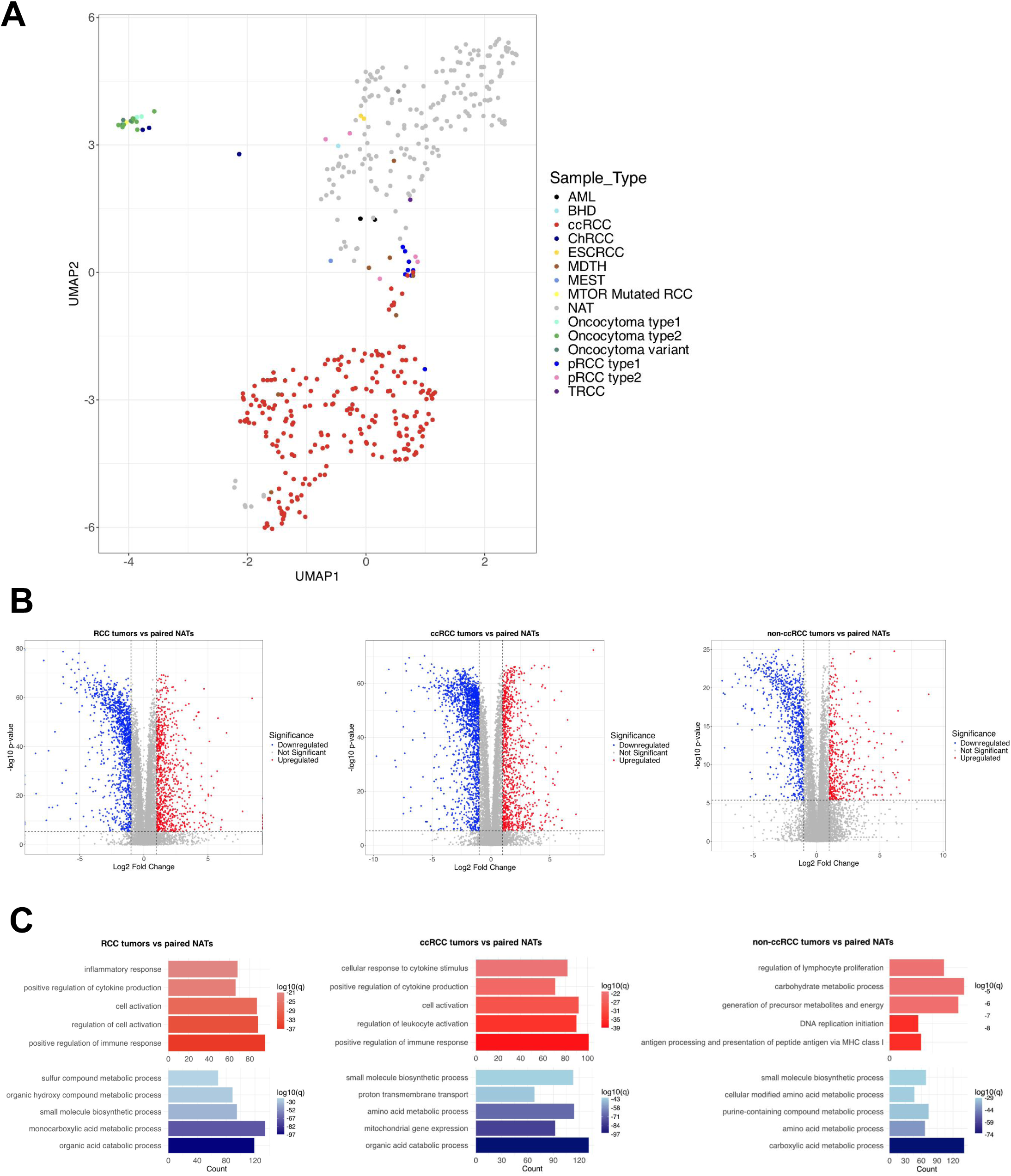
A. Uniform manifold approximation and projection analysis of the RCC tumors and paired NATs. B. Differential analysis between RCC tumors *vs* paired NATs (left), ccRCC tumors *vs* paired NATs (middle), and non-ccRCC tumors *vs* paired NATs (right). Significantly altered proteins were defined as > 2-fold changes with a Bonferroni adjusted *p*< 0.05. C. Analysis of significantly differentially regulated pathways (adjusted *p* < 0.05) between RCC tumors *vs* paired NATs (left), ccRCC tumors *vs* paired NATs (middle), and non-ccRCC tumors *vs* paired NATs (right). Red bars indicated pathways that were upregulated in tumors, and blue bars indicate pathways that were downregulated in tumors.

To discover the differences between RCC tumors and NATs, we used Wilcoxon Rank Sum test to compare the differentially expressed proteins (DEPs) for the following: RCC tumors vs paired NATs, ccRCC tumors vs paired NATs, and non-ccRCC tumors vs paired NATs. Compared to the paired NATs, 836 and 1166 proteins were upregulated and downregulated in RCC tumors, respectively(Figure 1B and Table S1). The comparison between the ccRCC tumors and paired NATs showed that 2495 proteins significantly changed with 910 proteins upregulated and 1585 downregulated in the ccRCC tumors relative to the paired NATs (Figure 1B and Table S1). For the comparison between the non-ccRCC tumors and the paired NATs, the results showed 1262 proteins significantly changed with 459 proteins upregulated and 803 downregulated in the non-ccRCC tumors relative to the paired NATs (Figure 1B and Table S1). Enrichment analysis revealed positive regulation of immune response, cell activation, and positive regulation of cytokine production to be upregulated in RCC tumors, and organic acid catabolic process and small molecule biosynthetic process to be downregulated (Bonferroni adjusted *p* < 0.05, Figure 1C and Table S1). Similar results were found in ccRCC tumors compared with the NATs (Figure 1C and Table S1). For the comparison between non-ccRCC tumors and NATs, the enrichment analysis revealed DNA replication initiation, antigen processing and presentation of peptide antigen via MHC class Ⅰ, and generation of precursor metabolites and energy to be upregulated in non-ccRCC tumors, and carboxylic acid metabolic process and purine-containing compound metabolic process to be downregulated (Bonferroni adjusted *p* < 0.05, Figure 1C and Table S1). These pathways were not enriched as top 5 pathways for both RCC tumors and ccRCC tumors when compared with the NATs (Figure 1C). Of note, the number of ccRCC tumors accounts for most of the RCC tumors (Figure S1A), thus, the difference between non-ccRCC tumors and NAT was obscured.

### Systematic sample selection of the ccRCC tumors

To fully capture the complexity and diversity of RCC tumors for both ccRCC and non-ccRCC, we selected representative ccRCC samples for our study. The selection was guided by the following steps: First, gene mutation-based sample selection was performed to represent the diversity of ccRCC tumors at the genetic level. We focused on mutation profiles in key genes known to be involved in ccRCC tumors, namely VHL, SETD2, PBRM1, KDM5C, and BAP1. We selected at least four samples for each mutation site (Figure S2A and Table S2). This approach ensured a broad representation of the genetic heterogeneity inherent in ccRCC tumors. Our second step considered the diversity of histopathological, immune, methylation, and NMF subtypes which were established in our previous study^24^ for ccRCC tumors. We selected samples representing each of the four histopathological subtypes (CL, CH, CH-S, CH-R), four immune subtypes (CD8+ inflamed, CD8- inflamed, Metabolic desert, VEGF desert), three methylation subtypes (Methyl1, Methyl2, Methyl3), and three NMF subtypes (NMF1, NMF2, NMF3) (Figure S2B and Table S2). This allowed us to capture the biological and molecular diversity in ccRCC tumors as representative ccRCC tumors for the RCC cohort. The third step was to consider the patient’s gender, grade, and stage to ensure that the selected samples were representative of these clinical characteristics (Figure S2C and Table S2). This meant incorporating a balanced mix of male and female patients’ samples, thus accounting for potential gender-specific variations in ccRCC tumors. We also selected samples across different tumor grades, including low-grade (G1 and G2) and high-grade (G3 and G4) tumors (Figure S2C and Table S2). This was crucial to capture the proteomic differences associated with tumor aggressiveness and potential variations in disease progression. In addition, we considered the stage of the disease at the time of sample collection. Our selection included samples from early (Stages I and II) to advanced stages (Stages III and IV) of ccRCC (Figure S2C and Table S2). This allowed us to account for the progression-related changes in the proteomic profiles of ccRCC tumors and understand how these changes might influence disease outcome. Then, we verified that the proteomic data for our selected samples encompassed all the protein groups identified in the proteomic data of ccRCC tumors (Table S2). Finally, we profiled the phenotypes of selected ccRCC samples (Figure 2A and Table S2). By systematically selecting samples that accurately represent the diversity of ccRCC tumors in terms of their genetic difference along with histopathological, immune, methylation, NMF subtypes, and clinical characteristics (gender, grade, and stage), we believe we have captured a comprehensive snapshot of the complex and heterogeneous nature of ccRCC tumors. This will allowed us to make a more detailed and comprehensive interpretation of RCC in subsequent analyses, thereby increasing the possibility of making new discoveries.

**Figure 2.**
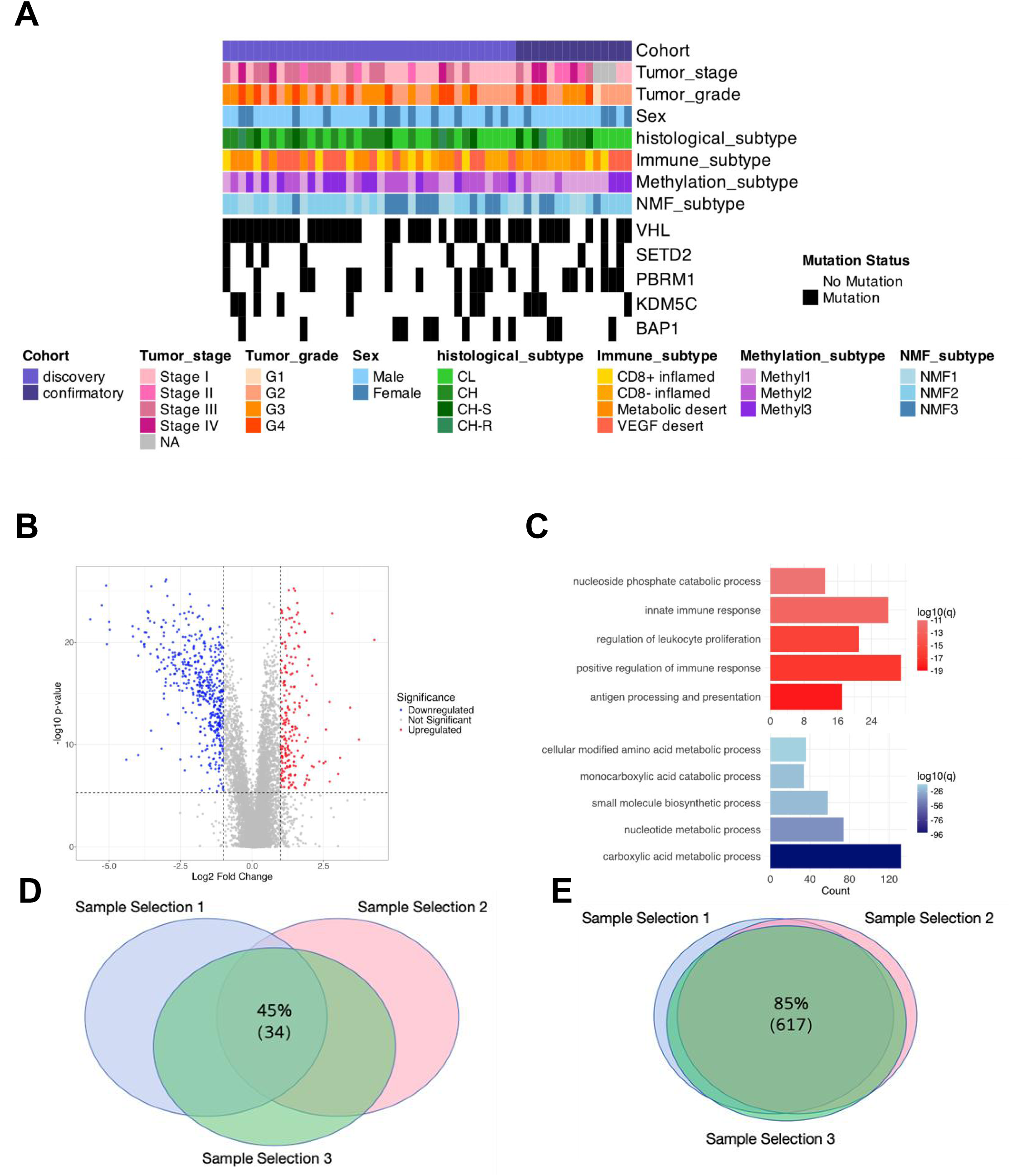
A. The clinical phenotypes profiling of proteomic data of the selected 53 ccRCC tumors. B. Differential analysis between the selected RCC tumors vs paired NATs. Significantly altered proteins were defined as > 2-fold changes with a Bonferroni adjusted *p*< 0.05. C. Analysis of significantly differentially regulated pathways (adjusted *p* < 0.05) between the selected RCC tumors vs NATs. Red bars indicated pathways that were upregulated in tumor tissues, and blue bars indicated pathways that were downregulated in tumor tissues. D. Venn diagram for the sample ID overlap between 3 times sample selections. E. Venn diagram for the DEPs overlap between 3 times sample selections.

### Proteomic alterations of RCC tumors compared to NATs

To fully understand the differences between RCC tumors and NATs, we compared protein expressions between RCC tumors composed of selected representative ccRCC tumors (Figure 2A and S2) and non-ccRCC tumors and paired NATs. In total, 681 proteins showed significant differential expressions with 209 proteins upregulated and 472 downregulated in RCC tumors compared to NATs (Figure 2B and Table S2). Enrichment analysis revealed differential expressions of proteins involved in various biological pathways between the RCC and NAT samples. Specifically, we identified antigen processing and presentation, positive regulation of immune response, and regulation of leukocyte proliferation that were upregulated, carboxylic acid metabolic process, nucleotide metabolic process, and small molecule biosynthetic process were downregulated in the RCC tumors compared to the NATs (Figure 2C and Table S2). The upregulation of proteins suggested an active immune response in RCC tumors. On the other hand, the downregulation of proteins indicated a potential reprogramming of metabolic pathways in RCC tumors, which might contribute to cancer cell survival and growth. The differentially expressed proteins and the associated biological pathways identified in our study provide valuable insights into the molecular mechanisms underlying RCC oncogenesis and progression. To validate the representativeness of the differential proteins identified in our analysis of the selected RCC and NAT samples, we implemented the same sample selection strategy 3 times and 45% of the samples overlapped (Figure 2D). Next, we compared the differential proteins identified in each selection. Remarkably, there was a consistent overlap of 85% in the differential proteins identified across all three groups (Figure 2E). This suggested a strong representativeness and reliability of our sample selection approach.

### Protein signature identification for RCC tumors

While the differential analysis of representative samples illustrated the differential proteins between RCC and NAT, reflecting common distinguishing features among various RCC subtype samples against NAT, this analysis did not represent the individual characteristics of each RCC or NAT sample. Furthermore, the vast number of differential proteins made the discovery of the most important protein signatures that could distinguish RCC from NAT challenges. To identify the protein signatures from the proteomic data between RCC tumors and NATs, a comprehensive ML exercise including feature selection and permutation validation was carried out on the selected RCC dataset with selected ccRCC tumors, non-ccRCC tumors, and paired NATs. For the feature selection, a Random Forest classifier with Recursive Feature Elimination and 5-fold Cross-Validation (RFECV) was applied to the proteins using 20% of samples from the selected RCC dataset. To robustly train and evaluate the model, the 5-fold cross-validation process divided the data into five subsets, training the model iteratively on four folds while testing it on the fifth. Accuracy scores were calculated for each fold, and the mean and standard deviation of accuracies were recorded to assess the model’s consistency across folds. After model training, RFECV optimized feature selection by iteratively removing the least important features based on the Random Forest model’s feature importance scores. RFECV selected only the most relevant proteins by evaluating feature subsets through cross-validation, minimizing model complexity while preserving accuracy. RFECV resulted in selecting 55 protein signatures from the selected RCC dataset in segregating RCC tumors from NATs (Table S3). The heatmap illustrated the differences between the RCC tumors and NATs, as well as the similarities between ccRCC and non-ccRCC (Figure 3A). After feature selection, the permuted dataset with randomly shuffled RCC tumor and NAT labels was used to assess whether the original model performance was better than random chance. The higher receiver operating characteristic (ROC) curve and area under the curve (AUC) observed with the original labels compared to the permuted labels confirms the predictive value of these protein signatures in differentiating between RCC tumor and NAT samples (Figure 3B). The 55 protein signatures selected by RFECV included proteins with particularly high importance scores (Table S3). Among these, the top 3 proteins further confirmed their key role in differentiating RCC tumors from NAT samples with high AUCs (Figure S3A-C). In addition, although the sample selection balanced the number of samples between ccRCC and non-ccRCC, the removal of ccRCC tumor and NAT samples may lead to insufficient representation of the protein signatures. To evaluate the representative of the 55 selected protein signatures, the same strategies were used to build an ML model with all patient samples from the entire RCC dataset (Figure S3D). The ROC curve and AUC confirmed the predictive value of these protein signatures in distinguishing RCC tumors from NATs in the entire RCC dataset. Notably, 28 of the 55 protein signatures identified by the ML model overlapped with the DEPs (Figure S3E). Through an ML approach, including feature selection and permutation validation, we identified 55 protein signatures from proteomic data that distinguish between RCC tumors and NATs.

**Figure 3.**
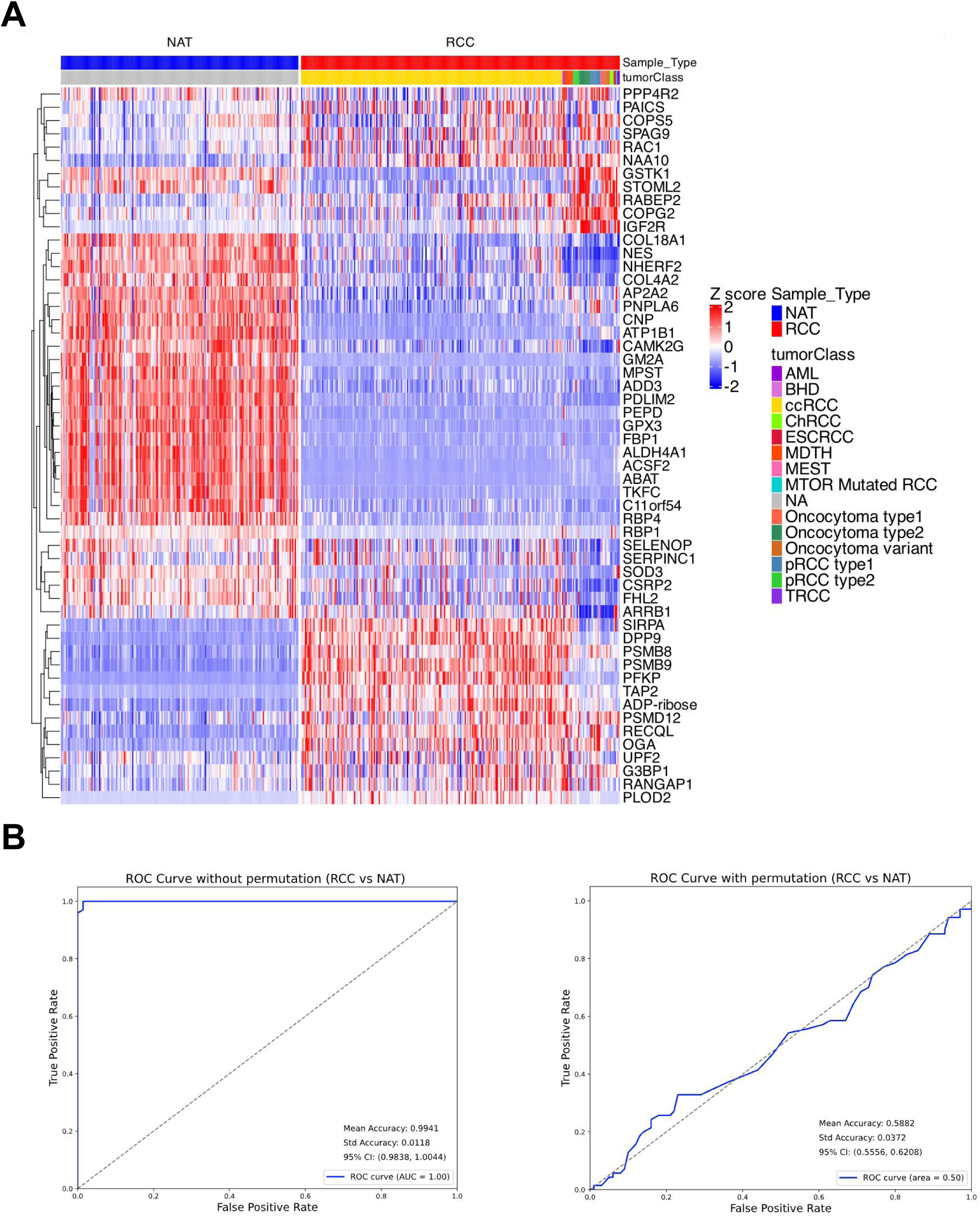
A. Heatmap representation of the protein signatures for the RCC tumors. B. The receiver operating graph of the protein signatures selected by the Random Forest classifier on the selected RCC tumors and NATs hold-out test dataset with (right) or without permutation (left). The area under the curve (AUC) was calculated.

### Protein signature identification for the RCC tumors subtypes

Following the identification of the protein signatures that distinguish RCC tumors from NATs, we endeavored to discern the proteomic disparities among various RCC subtypes. To establish the protein signatures for RCC subtypes, an ML exercise was performed on the selected RCC dataset without the NATs. RFECV was applied to the data, using 20% of patient samples for feature selection. This process resulted in the selection of 39 protein signatures that distinguished between RCC tumor subtypes (ccRCC tumors, Oncocytomas, pRCC tumors, and other non-ccRCC tumors, Table S3). The heatmap showed the protein signatures for the RCC tumor subtypes (Figure 4A). To validate these protein signatures, a permuted dataset with shuffled RCC tumor subtype labels was used. The model’s higher performance with the original labels compared to the permuted labels confirmed the predictive value of these protein signatures (Figure 4B). While individual proteins perform well in distinguishing between RCC and NAT (Figure S3A-C), the top 3 important proteins within the set of 39 protein signatures (Table S3) do not exhibit a satisfactory performance in distinguishing between RCC tumor subtypes (Figure S4A-C). Additionally, an ML model was built with samples from the entire RCC tumors to further evaluate the representative of the 39 selected protein signatures. The receiver operating graph confirmed the predictive value of these protein signatures in distinguishing RCC tumor subtypes in the entire RCC dataset (Figure S4D). Interestingly, the protein PNPLA6 overlaps between the protein signatures distinguishing RCC from NAT and those distinguishing different RCC tumor subtypes (Figure S4E). It is downregulated in RCC compared to NAT, while being upregulated in pRCC compared to other RCC tumor subtypes (Figure S4F and G). However, the ROC curve indicates that this protein alone does not effectively differentiate between RCC or NAT, nor among different RCC subtypes (Figure S4H and I). Considering our previous finding that the top 3 important proteins in the protein signatures were insufficient in distinguishing different RCC tumor subtypes, it seems challenging to rely on a single protein for differentiation. Instead, a combination of multiple proteins should be considered for a more accurate characterization. Through the ML approach, we identified 39 protein signatures that distinguish between different RCC tumor subtypes, including ccRCC tumors, Oncocytomas, pRCC tumors, and other non-ccRCC tumors.

**Figure 4.**
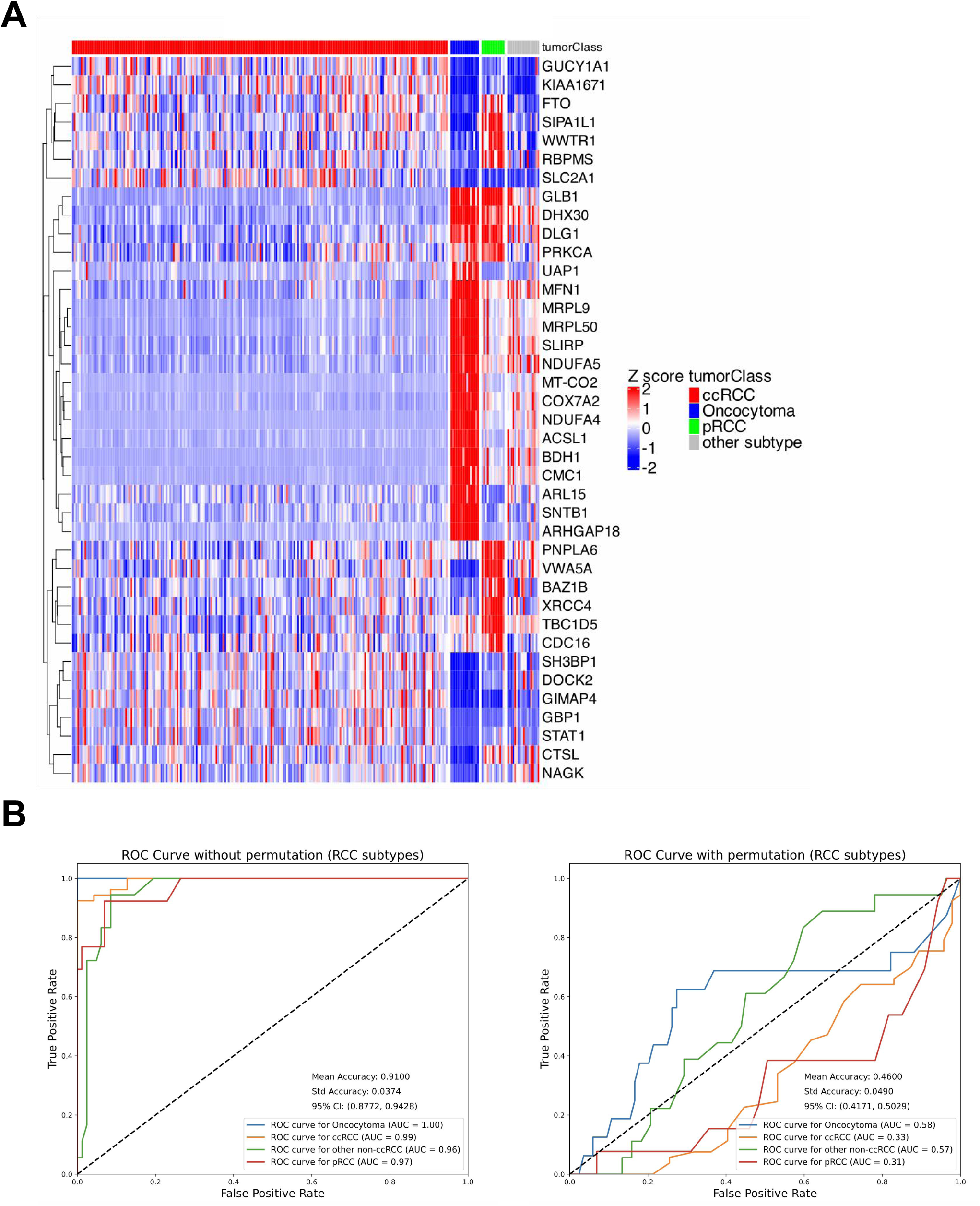
A. Heatmap representation of the protein signatures for the RCC tumor subtypes (ccRCC tumors, oncocytomas, PRCC tumors, and other non-ccRCC tumors). B. The receiver operating graph of the protein signatures selected by the classifier Random Forest on the selected ccRCC tumors, oncocytoma, PRCC tumors, and other non-ccRCC tumors hold-out test dataset with (right) or without permutation (left). The AUC was calculated.

## Discussion

In this study, DIA-based proteomics data provides high-quality data sources, which can be further investigated to gain deeper insights into disease biology. Our study presented an approach to select representative ccRCC samples to be analyzed with other non-ccRCC subtypes to capture the diversities of RCC. The analysis of the DIA proteomic data from a broad range of samples highlighted the common and unique characteristics of ccRCC tumors relative to non-ccRCC tumor subtypes. We carried out ML analyses to identify protein signatures that can distinguish ccRCC tumor subtypes and RCC tumors from NATs. These protein signatures were validated by permutation and across the entire RCC dataset.

Understanding the molecular differences between RCC tumors and normal tissues is crucial for improving diagnostic accuracy and treatment strategies. The UMAP analysis using proteomic data revealed different clusters of RCC tumors. Specifically, we identified the Oncocytoma type 1, Oncocytoma type, and Oncocytoma variant distinct from ccRCC tumors and NATs (Figure 1A). This result was consistent with the significant genome difference between the Oncocytomas and other RCC subtypes^40^. This further suggested that proteomic profiling could potentially aid in distinguishing Oncocytomas from malignant RCC subtypes, addressing a challenge in RCC diagnosis and management^41–43^. This could potentially lead to improved diagnostic accuracy and better patient management in RCC.

An important aspect of our study was the comparison of protein expressions between RCC tumors and paired NATs. However, the predominance of ccRCC samples over non-ccRCC samples in the RCC posed a challenge, as the characteristics of non-ccRCC were overshadowed by those of ccRCC (Figure 1B, Figure 1C, and Figure S1A). To address this issue, we selected a subset of ccRCC samples based on molecular pathology features representative of ccRCC, ensuring their number was balanced with that of non-ccRCC samples. Our approach to ccRCC tumor sample selection included consideration of genetic mutation sites and different histopathological, immune, methylation, and NMF subtypes as well as stage and grade, thus capturing the biological and molecular diversity in ccRCC. After completing three separate rounds of sample selection, we found the DEPs between RCC and NAT for each selection. The results showed that 85% of the DEPs were consistently identified across all three comparisons (Figure 2D and E). This approach emphasized the point of balanced samples with representative ones when studying heterogeneous diseases. Additionally, this sample selection process demonstrated the feasibility of selecting representative samples based on known molecular subtypes within large cohorts. The DEPs revealed differentially expressed pathways, antigen processing and presentation, positive regulation of immune response, and regulation of leukocyte proliferation were up-regulated while the carboxylic acid metabolic process, nucleotide metabolic process, and small molecule biosynthetic process were down-regulated in RCC tumors compared to NATs (Figure 2C). The upregulated proteins in RCC tumors aligned with earlier studies highlighting the role of the immune system in cancer progression^44,45^. On the other hand, the downregulation of proteins involved in metabolic processes resonated with the work of Hakimi et al. who demonstrated the reprogramming of metabolic pathways in ccRCC tumors and its potential role in promoting tumor growth and survival^46^. Furthermore, our results were in line with the study by Clark et al., where they found differential pathway changes in ccRCC tumors compared to NATs, indicating the profound molecular alterations that occur during RCC carcinogenesis^25^. It suggested that despite the heterogeneity within RCC tumors, there were common protein expression patterns that can be reliably identified.

Building on these findings, our study utilized an ML approach, including feature selection and permutation validation, to identify protein signatures in proteomic data that could differentiate RCC tumors from NATs, as well as distinguish between different RCC tumor subtypes. Our ML exercise, which employed a Random Forest classifier with RFECV, identified 55 protein signatures that distinguished RCC tumors from NATs (Figure 3A). Among these proteins, several had been reported to be related to RCC. Wang et al. found that Ras GTPase-activating protein-binding protein 1 (G3BP1) was significantly higher in RCC tumors comparing to NATs, and knockdown of G3BP1 decreased tumor cell growth and metastasis^47^. Liu et al. reported that reduced glutathione peroxidase 3 (GPX3) in primary ccRCC due to promoter methylation was associated with a poor prognosis^48^. Studies have shown that loss of fructose-1,6-bisphosphatase 1 (FBP1) expression was a hallmark of ccRCC and contributes to the metabolic reprogramming of cancer cells (known as the Warburg effect)^49,50,50,51^. Reduced FBP1 levels are associated with tumor growth and poor prognosis^51^. In line with these findings, our study also observed a significant downregulation of FBP1 protein levels in RCCs when compared to NATs (Table S1). This consistent observation strengthens the association of FBP1’s role in the pathology of RCC. Building on recent research by Liu et al., which reported an upregulation of PLOD2 under hypoxic conditions in ccRCC and associated high PLOD2 expression with poor prognosis in ccRCC patients^52^. We observed a significant increase in PLOD2 levels in RCCs compared to NATs (Table S1), thus lending further support to the potential role of PLOD2 in the pathology of ccRCC. These proteins have been identified through various studies as having significant roles in the development, progression, or prognosis of RCC tumors. To be noticed, 28 of the 55 protein signatures identified by the ML model overlapped with the DEPs (Figure S3E). The primary advantage of the ML approach lies in its capacity to handle high-dimensional data and consider intricate relationships between variables. By identifying 55 proteins, ML possibly recognized complex patterns and interactions among these proteins that may not be evident when considering each protein individually. This smaller set may be more biologically relevant, potentially reflecting key pathways or processes intrinsic to RCC pathogenesis. On the other hand, the Wilcoxon test did not consider potential interactions among proteins which provided a broad view of the differential proteomic landscape between RCC and NATs. The overlap of 28 proteins between the two methods provided a subset of proteins that are both statistically significant and potentially part of the complex biological interactions relevant to RCC.

The distinction between ccRCC and non-ccRCC tumors underscored the need for more specific protein signatures that can accurately distinguish between RCC tumor subtypes (Figure 3A). In response to this challenge, we used a ML approach and successfully identified 39 protein signatures that differentiate among RCC tumor subtypes. These subtypes include ccRCC tumors, Oncocytomas, pRCC tumors, and other non-ccRCC tumors (Figure 4A). Notably, within these identified protein signatures, several proteins have already been reported to have associations with either non-ccRCC or ccRCC, further validating the relevance of our findings. For instance, the solute carrier family 2, facilitated glucose transporter member 1 (SLC2A1) shows differential expression in various RCC subtypes, with high expression in ccRCC and low expression in non-ccRCC subtypes such as pRCC^53,54^. Additionally, mitochondrial proteins such as Mitofusin-1 (MFN1), Cytochrome c oxidase subunit NDUFA4 (NDUFA4), and NADH dehydrogenase [ubiquinone] 1 alpha subcomplex subunit 5 (NDUFA5), implicated in mitochondrial dynamics and complex I function respectively, may be associated with the pathological features of Oncocytomas, characterized by mitochondrial accumulation^55,56^. Interestingly, we found that the protein PNPLA6 emerged in both the protein signature set distinguishing RCC tumors from NATs and the protein signature set differentiating RCC tumor subtypes (Figure S4E). The expression of PNPLA6 was lower in RCC tumors compared to NATs, and among the various RCC subtypes, its expression was highest in pRCC compared to other subtypes (Figure S4F and G). Nonetheless, the ROC-AUC indicated that this protein alone was not sufficient to effectively distinguish RCC tumors from NATs or among different RCC tumor subtypes. However, in differentiating RCC tumor subtypes, it showed some utility in distinguishing pRCC from other subtypes (Figure S4H and I). We propose that accurately distinguishing among RCC tumor subtypes using a single protein is challenging due to the high heterogeneity of RCC tumor subtypes. Conversely, distinguishing RCC tumors from NATs using a single protein showed relatively better results, suggesting a certain level of internal similarity within RCC. However, given the heterogeneity among RCC tumor subtypes, a combination of multiple proteins is likely required to accurately distinguish RCC tumors from NATs, as well as to differentiate among the various RCC tumor subtypes. In summary, our ML approach has facilitated the identification of protein signatures that differentiate among RCC tumor subtypes, contributing to a refined understanding of RCC pathology.

We noticed the limitation that the number of the Oncocytoma and pRCC samples in our study cohort was much larger than that of the other subtypes of non-ccRCC samples such as AML and BHD which had only 1 or 2 samples. This imbalance may have obscured some unique characteristics of non-ccRCC tumors. Future studies could benefit from expanding the non-ccRCC sample size to provide a more balanced comparison. The identified protein signatures could contribute to personalized treatment strategies. However, further validation studies are needed to confirm the predictive value of these signatures.

In conclusion, we developed a sample selection approach to balance the sample number between ccRCC tumors and non-ccRCC tumors and considered a variety of factors to choose representative samples. We used ML to find protein signatures that could differentiate RCC tumors from NATs and differentiate between various RCC tumor subtypes. The similarities and differences between the different RCC tumor subtypes were emphasized by these protein signatures. Ultimately, this study offered a comprehensive DIA-based proteomics data source for RCC, which is a helpful resource for further research.

### Experimental Model and Subject Details

#### MS sample processing and data Collection

In this study, we performed proteomics profiling of 48 non-ccRCC tumors^24–26^ and 213 ccRCC tumors^24,26^. The mutation sites data was available for 261 tumor samples^24–26^. The 48 non-ccRCC samples had been described previously^26^. There are 2 ccRCC tumor samples with labeled C3L-00908-T-1 and C3L-00908-T-2, which were from different aliquots of the same case ID C3L-00908-T. In the subsequent data analysis, C3L-00908-T-1 was used and C3L-00908-T-2 was removed.

#### Sample processing for protein extraction and tryptic digestion

All samples for the current study were prospectively collected as described above and processed for MS analysis, tissue lysis and downstream sample preparation for proteomic analysis were carried out as previously described^24–26^.

#### EvoSep-timsTOF for proteomic analysis

All the LC-MS/MS data were acquired via EvoSep coupled with timsTOF HT (Bruker) in data-independent acquisition mode. The methods for acquiring proteomics were described previously^57^.

#### MS data analysis

The spectral library was created using Spectronaut® 18.4 (Biognosys AG) by merging all search archives from both RCC and NAT samples. The mass tolerance of MS and MS/MS was dynamically set with a correction factor of one in the search settings. All raw files were matched against a unified Homo sapiens GENCODE42 protein sequence database, which had an equal number of decoy sequences appended. We applied a Q value cutoff of 0.01 for precursor filtering, corresponding to an FDR of 1%. A fixed modification was set for Carbamidomethyl (C) while Acetyl (Protein N-term) and Oxidation (M) were determined as variable modifications. The peptide quantification was derived from the sum of the quantities of its top 3 precursors. Meanwhile, precursor quantity was calculated by taking the total area of its top 3 fragment ions at the MS/MS level. The data was normalized by being divided by the median of each sample. Differential analysis was carried out by calculating the mean log2 fold changes between RCC tumors vs. paired NATs, ccRCC tumors vs. paired NATs, non-ccRCC tumors vs. paired NATs, selected RCC tumors vs. paired NATs. A Wilcoxon Rank Sum Test was performed on each protein to compare the median expression levels between two independent groups. Proteins with an adjusted p-value below a Bonferroni-corrected threshold were considered significantly different. Alongside the statistical test, log2 fold changes were calculated to determine the direction and magnitude of expression differences, classifying proteins as “Upregulated” or “Downregulated”.

#### Functional enrichment analysis

Ontology enrichment analysis of the DEPs was conducted using the metascape^58^ available at https://metascape.org with default settings. Supplementary Table 2 includes the list of significantly enriched pathway terms^59,60^ and associated proteins. The gene ontologies were considered for biological processes.

#### Machine learning model construction

Three steps, from feature selection and feature significance validation to model performance evaluation, were included in the ML framework for this study. An RCC protein matrix was utilized as the input, with each row representing a protein and each column representing an RCC sample involved in the task. To select protein signatures that distinguish RCC samples, a random forest (RF) classifier with Recursive Feature Elimination and 5-fold Cross-Validation (RFECV) was applied. RFECV is a method for feature selection that iteratively fits a model and removes the least important features based on their impact on model performance, with each iteration validated through 5-fold cross-validation. It helps in determining the smallest number of features that yield the maximum predictive power, which is crucial for model simplicity and interpretability. Initially, 5-fold cross-validation was defined using sklearn.model_selection.StratifiedKFold. Then RFECV was executed using sklearn.feature_selection.RFECV, employing RF with sklearn.ensembl.Random ForestClassifer as the classifier with default parameters. After feature selection, a permutation test was conducted to validate the significance of selected features. This involved randomly shuffling the labels while maintaining their original proportions and then re-training the model with these permuted labels using the same set of features initially selected. The model’s performance was then evaluated using the ROC-AUC metric. A comparison of the ROC-AUC scores between the model trained with original labels and the model trained with permuted labels showed that the original labels yielded significantly higher scores. This confirmed that the selected features possess predictive value and are not capturing patterns due to random chance, thus validating their importance in accurate classification.

## Data availability

The datasets during and/or analyzed during the current study are available from the corresponding author on reasonable request.

## ACKNOWLEDGMENTS

This work was supported by the National Institutes of Health, National Cancer Institute, Clinical Proteomic Tumor Analysis Consortium (CPTAC, U24CA271079).

---

**Figure S1.**
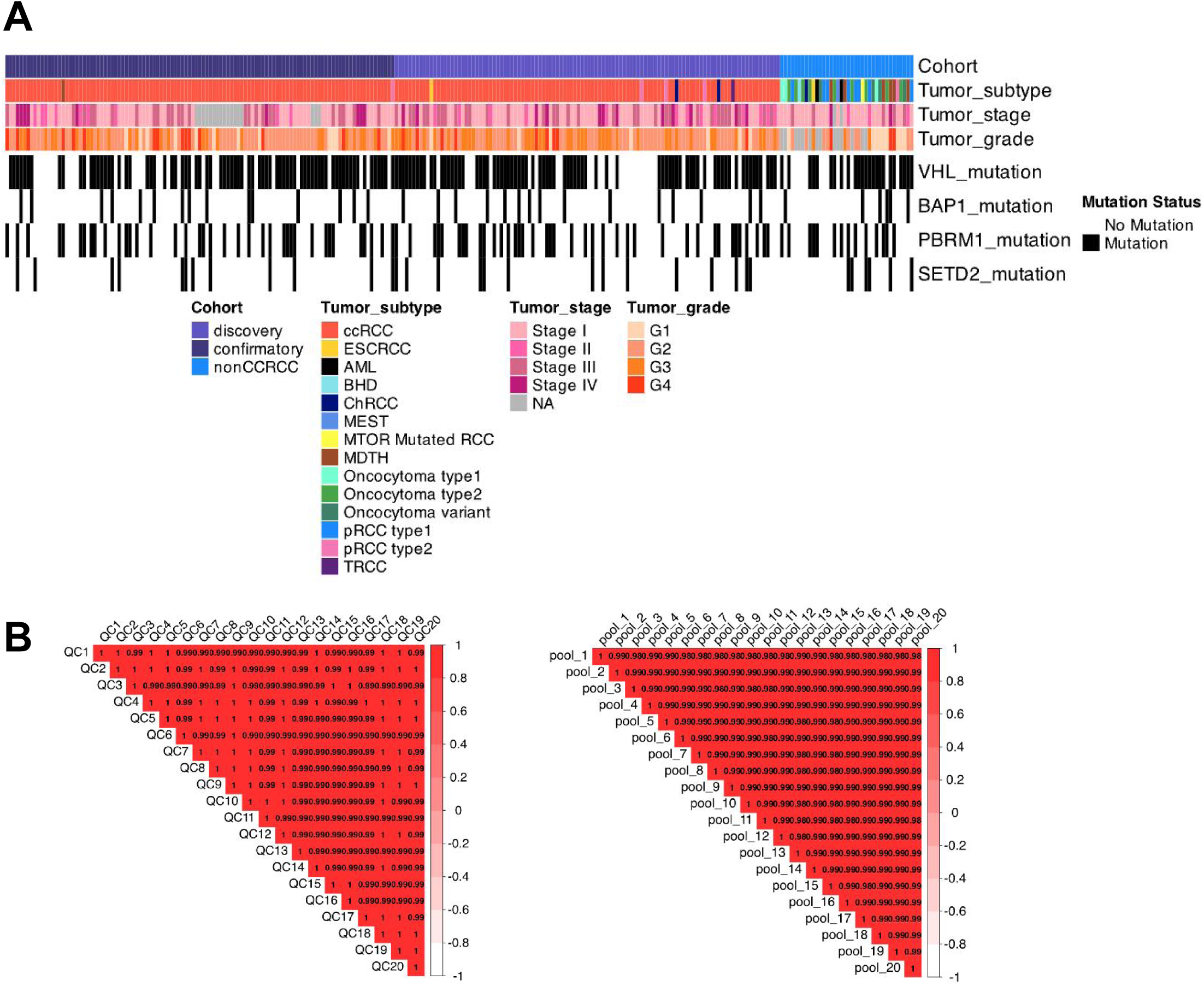
**A.** The clinical phenotypes profiling of the RCC tumors. **B.** Correlation analysis of 20 NCI-7 QC samples (left) and 20 pool samples (right) respectively as MS quality control to evaluate the robustness of label-free quantification. The average correlation coefficient among the samples was 0.99.

**Figure S2.**
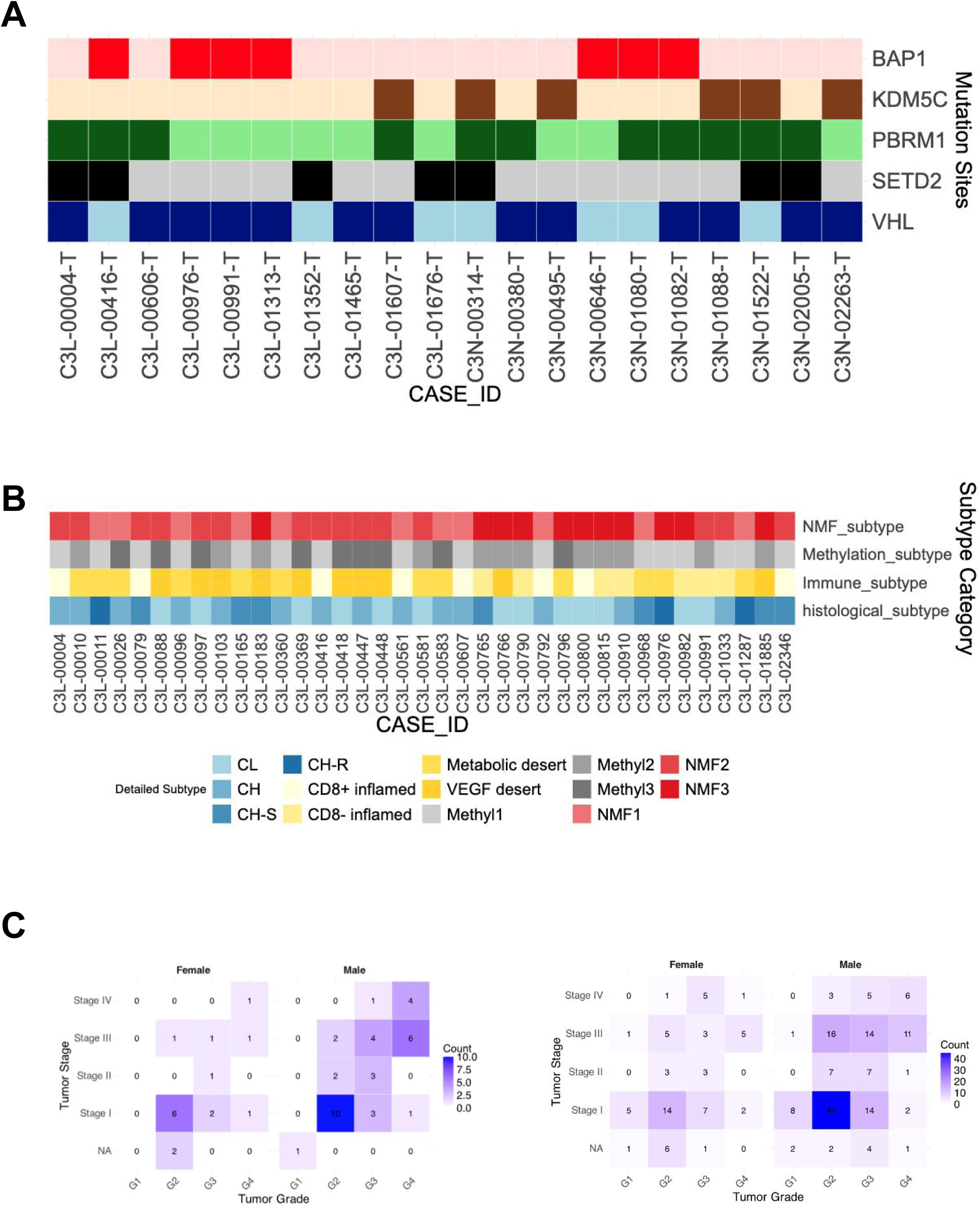
A. The mutation sites of the selected ccRCC tumors. B. The NMF, methylation, immune, histological subtypes of the selected ccRCC tumors. C. The number of ccRCC tumor patients in each grade, stage, and gender in the selected ccRCC samples (left) and in the entire ccRCC samples (right).

**Figure S3.**
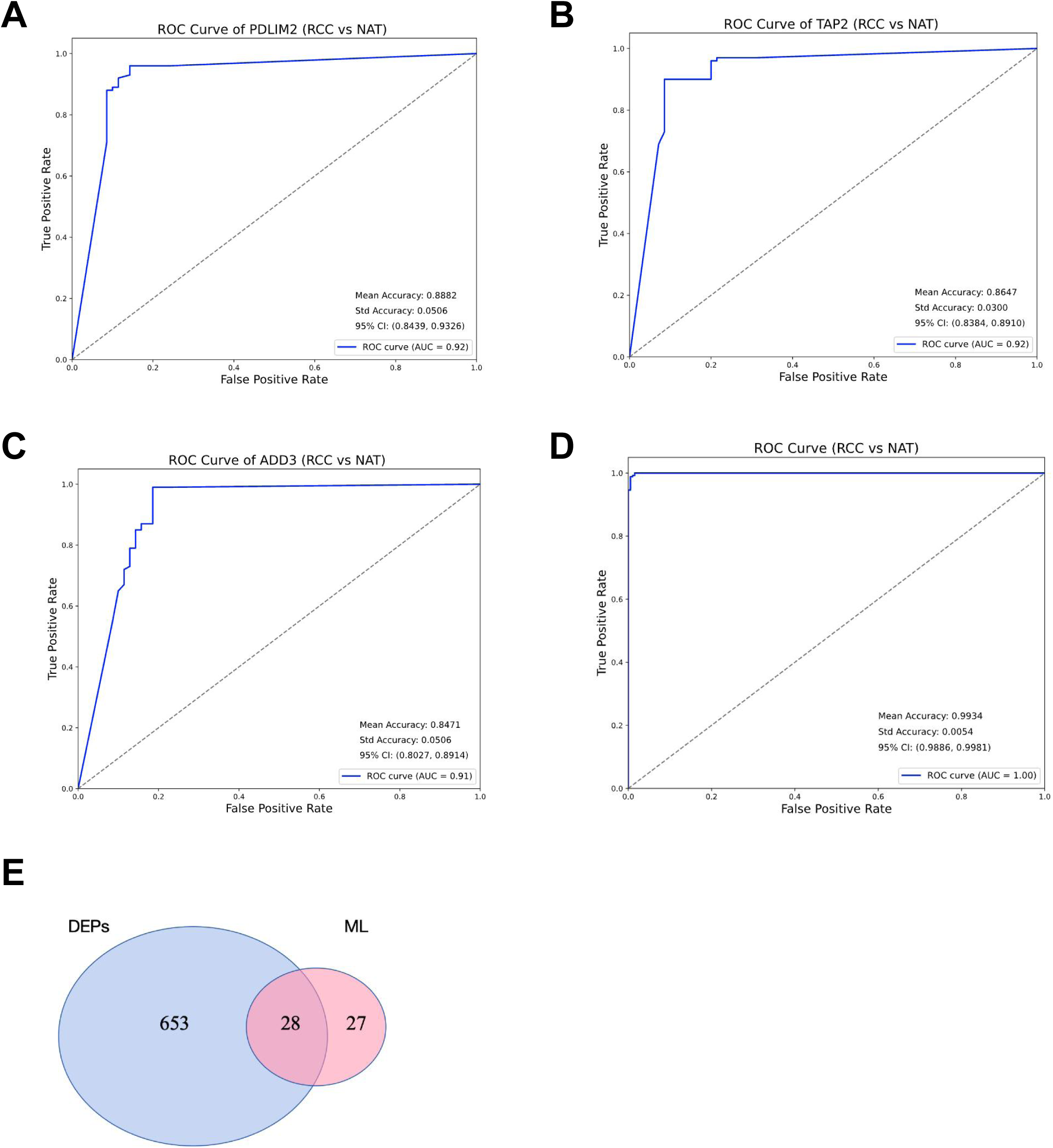
A-C. The receiver operating graph of the top 3 proteins in the protein signature list for the RCC tumors on the selected RCC tumors and NATs dataset. The AUC was calculated. D. The receiver operating graph of the protein signatures for the RCC tumors on the entire RCC tumors and NATs dataset. The AUC was calculated. E. Venn diagram for the protein signatures and the DEPs.

**Figure S4.**
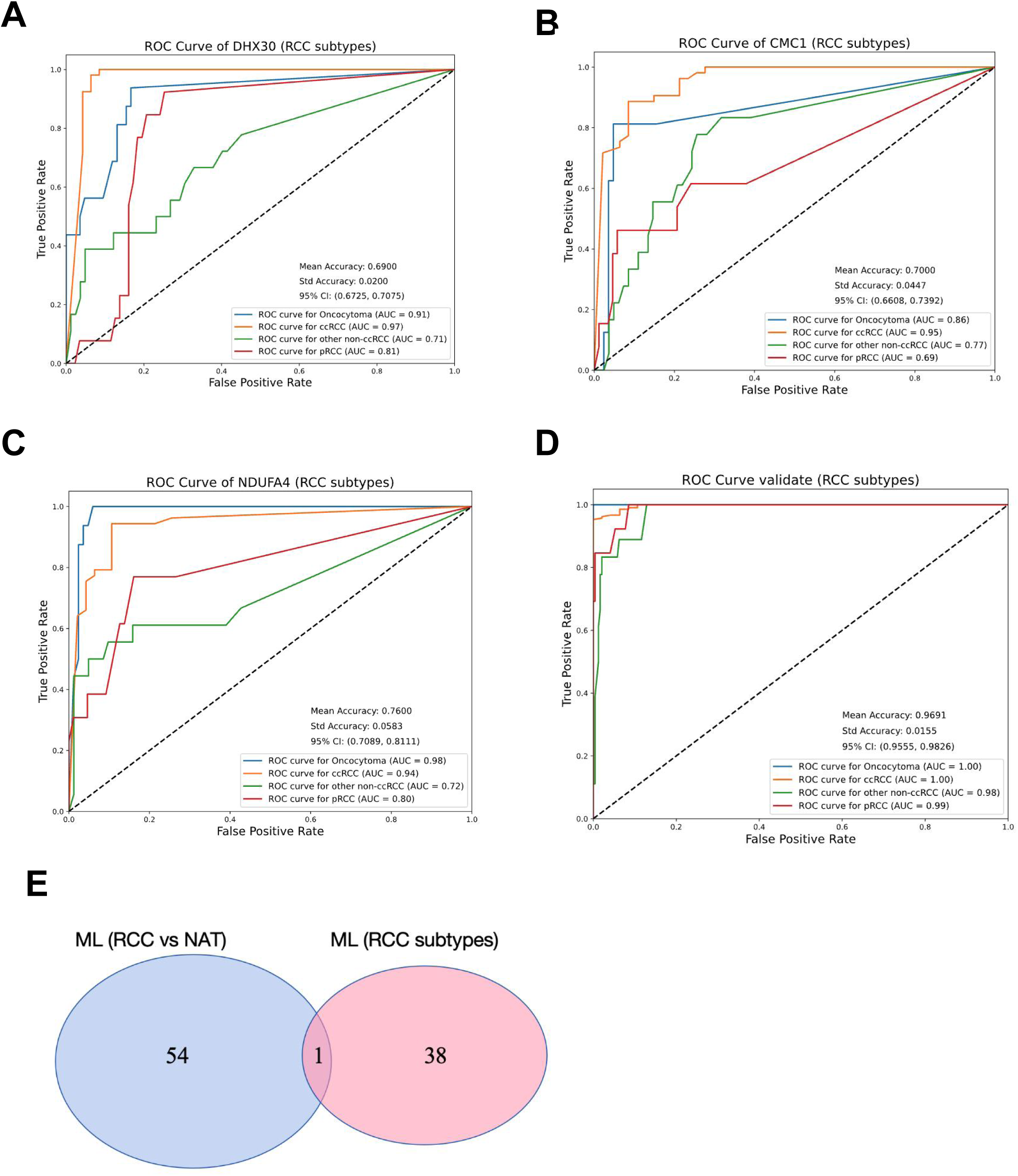
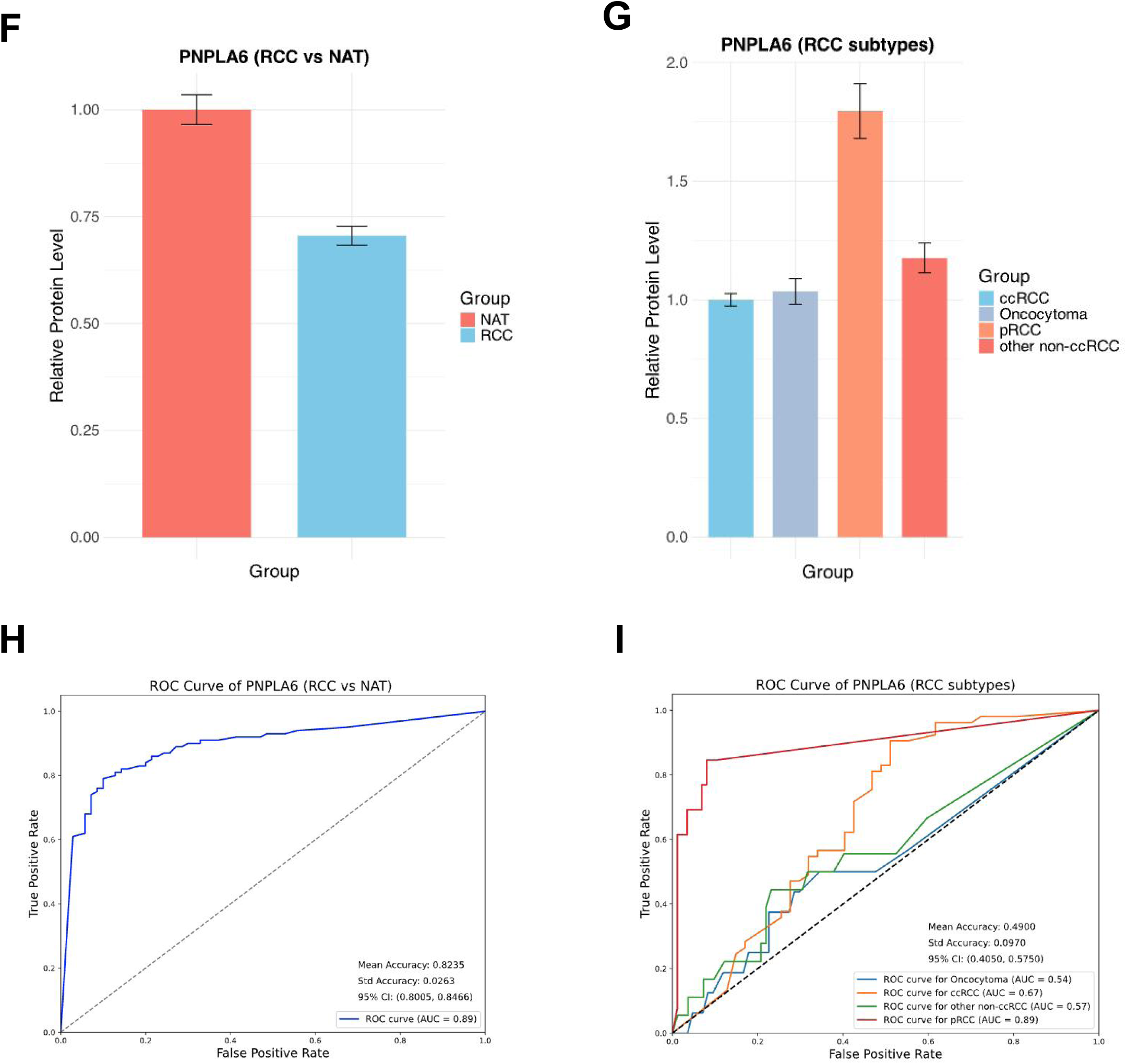
A-C. The receiver operating graph of the top 3 proteins in the protein signature list for the RCC tumor subtypes on the selected RCC tumors and NATs dataset. The AUC was calculated. D. The receiver operating graph of the protein signatures for the RCC tumor subtypes (ccRCC tumors, oncocytomas, PRCC tumors, and other non-ccRCC tumors) on the entire RCC tumors and NATs dataset. The AUC was calculated. E. Venn diagram for the protein signatures for RCC tumors and the protein signatures for RCC tumor subtypes. F. Bar plot of the relative protein level of PNPLA6 between the RCC tumors and NATs in the selected RCC tumor dataset. G. Bar plot of the relative protein level of PNPLA6 between the RCC subtypes in the selected RCC tumor dataset without the NATs. H. The receiver operating graph of PNPLA6 for the RCC tumors on the selected RCC tumor dataset. The AUC was calculated. I. The receiver operating graph of PNPLA6 for the RCC tumor subtypes on the selected RCC tumor dataset without the NATs. The AUC was calculated.

## Reference

1. Siegel, R. L., Giaquinto, A. N. & Jemal, A. Cancer statistics, 2024. CA: A Cancer Journal for Clinicians 74, 12–49 (2024).

2. Bray, F., et al. Global cancer statistics 2022: GLOBOCAN estimates of incidence and mortality worldwide for 36 cancers in 185 countries. CA: A Cancer Journal for Clinicians 74, 229–263 (2024).

3. Moch, H. et al. The 2022 World Health Organization Classification of Tumours of the Urinary System and Male Genital Organs—Part A: Renal, Penile, and Testicular Tumours. European Urology 82, 458–468 (2022).

4. Hsieh, J. J. et al. Renal cell carcinoma. Nature reviews. Disease primers 3, 17009 (2017).

5. Moch, H., Cubilla, A. L., Humphrey, P. A., Reuter, V. E. & Ulbright, T. M. The 2016 WHO Classification of Tumours of the Urinary System and Male Genital Organs—Part A: Renal, Penile, and Testicular Tumours. European Urology 70, 93–105 (2016).

6. Warren, A. Y. & Harrison, D. WHO/ISUP classification, grading and pathological staging of renal cell carcinoma: standards and controversies. World J Urol 36, 1913– 1926 (2018).

7. Creighton, C. J. et al. Comprehensive molecular characterization of clear cell renal cell carcinoma. Nature 499, 43–49 (2013).

8. Ricketts, C. J. et al. The Cancer Genome Atlas Comprehensive Molecular Characterization of Renal Cell Carcinoma. Cell Reports 23, 313–326.e5 (2018).

9. Hakimi, A. A., Pham, C. G. & Hsieh, J. J. A clear picture of renal cell carcinoma. Nat Genet 45, 849–850 (2013).

10. Kapur, P. et al. Effects on survival of BAP1 and PBRM1 mutations in sporadic clear-cell renal-cell carcinoma: a retrospective analysis with independent validation. The Lancet Oncology 14, 159–167 (2013).

11. Hakimi, A. A. et al. Adverse Outcomes in Clear Cell Renal Cell Carcinoma with Mutations of 3p21 Epigenetic Regulators BAP1 and SETD2: A Report by MSKCC and the KIRC TCGA Research Network. Clinical Cancer Research 19, 3259–3267 (2013).

12. Wang, X.-M. et al. TRIM63 is a sensitive and specific biomarker for MiT family aberration-associated renal cell carcinoma. Mod Pathol 34, 1596–1607 (2021).

13. Baba, M. et al. TFE3 Xp11.2 Translocation Renal Cell Carcinoma Mouse Model Reveals Novel Therapeutic Targets and Identifies GPNMB as a Diagnostic Marker for Human Disease. Molecular Cancer Research 17, 1613–1626 (2019).

14. Skala, S. L. et al. Next-generation RNA Sequencing–based Biomarker Characterization of Chromophobe Renal Cell Carcinoma and Related Oncocytic Neoplasms. European Urology 78, 63–74 (2020).

15. Wang, L. et al. VSTM2A Overexpression Is a Sensitive and Specific Biomarker for Mucinous Tubular and Spindle Cell Carcinoma (MTSCC) of the Kidney. The American Journal of Surgical Pathology 42, 1571 (2018).

16. Mertins, P. et al. Proteogenomics connects somatic mutations to signalling in breast cancer. Nature 534, 55–62 (2016).

17. Zhang, H. et al. Integrated Proteogenomic Characterization of Human High-Grade Serous Ovarian Cancer. Cell 166, 755–765 (2016).

18. Aebersold, R. & Mann, M. Mass-spectrometric exploration of proteome structure and function. Nature 537, 347–355 (2016).

19. Budayeva, H. G. & Kirkpatrick, D. S. Monitoring protein communities and their responses to therapeutics. Nat Rev Drug Discov 19, 414–426 (2020).

20. Zhang, B. et al. Proteogenomic characterization of human colon and rectal cancer. Nature 513, 382–387 (2014).

21. Lu, P., Vogel, C., Wang, R., Yao, X. & Marcotte, E. M. Absolute protein expression profiling estimates the relative contributions of transcriptional and translational regulation. Nat Biotechnol 25, 117–124 (2007).

22. Wang, D. Discrepancy between mRNA and protein abundance: Insight from information retrieval process in computers. Computational Biology and Chemistry 32, 462–468 (2008).

23. Wu, L. et al. Variation and genetic control of protein abundance in humans. Nature 499, 79–82 (2013).

24. Li, Y. et al. Histopathologic and proteogenomic heterogeneity reveals features of clear cell renal cell carcinoma aggressiveness. Cancer Cell 41, 139–163.e17 (2023).

25. Clark, D. J. et al. Integrated Proteogenomic Characterization of Clear Cell Renal Cell Carcinoma. Cell 179, 964–983.e31 (2019).

26. Li, G. X. et al. Comprehensive proteogenomic characterization of rare kidney tumors. Cell Reports Medicine 5, 101547 (2024).

27. Li, J., Smith, L. S. & Zhu, H.-J. Data-independent acquisition (DIA): An emerging proteomics technology for analysis of drug-metabolizing enzymes and transporters. Drug Discovery Today: Technologies 39, 49–56 (2021).

28. Krasny, L. & Huang, P. H. Data-independent acquisition mass spectrometry (DIA-MS) for proteomic applications in oncology. *Mol*. Omics 17, 29–42 (2021).

29. Li, K. W., Gonzalez-Lozano, M. A., Koopmans, F. & Smit, A. B. Recent Developments in Data Independent Acquisition (DIA) Mass Spectrometry: Application of Quantitative Analysis of the Brain Proteome. Front. Mol. Neurosci. 13, (2020).

30. Ludwig, C. et al. Data-independent acquisition-based SWATH-MS for quantitative proteomics: a tutorial. Molecular Systems Biology 14, e8126 (2018).

31. Bakalarski, C. E. et al. The Impact of Peptide Abundance and Dynamic Range on Stable-Isotope-Based Quantitative Proteomic Analyses. Journal of proteome research 7, 4756 (2008).

32. Fernández-Costa, C. et al. Impact of the Identification Strategy on the Reproducibility of the DDA and DIA Results. J. Proteome Res. 19, 3153–3161 (2020).

33. De Silva, S., Alli-Shaik, A. & Gunaratne, J. Machine Learning-Enhanced Extraction of Biomarkers for High-Grade Serous Ovarian Cancer from Proteomics Data. Sci Data 11, 685 (2024).

34. Mann, M., Kumar, C., Zeng, W.-F. & Strauss, M. T. Artificial intelligence for proteomics and biomarker discovery. cels 12, 759–770 (2021).

35. Reel, P. S., Reel, S., Pearson, E., Trucco, E. & Jefferson, E. Using machine learning approaches for multi-omics data analysis: A review. Biotechnology Advances 49, 107739 (2021).

36. Libbrecht, M. W. & Noble, W. S. Machine learning applications in genetics and genomics. Nat Rev Genet 16, 321–332 (2015).

37. Zou, J. et al. A primer on deep learning in genomics. Nat Genet 51, 12–18 (2019).

38. Chaudhary, K., Poirion, O. B., Lu, L. & Garmire, L. X. Deep Learning–Based Multi-Omics Integration Robustly Predicts Survival in Liver Cancer. Clinical Cancer Research 24, 1248–1259 (2018).

39. Osipov, A. et al. The Molecular Twin artificial-intelligence platform integrates multi-omic data to predict outcomes for pancreatic adenocarcinoma patients. Nat Cancer 5, 299–314 (2024).

40. Durinck, S. et al. Spectrum of diverse genomic alterations define non–clear cell renal carcinoma subtypes. Nat Genet 47, 13–21 (2015).

41. Mirkheshti, N. et al. Renal oncocytoma: a challenging diagnosis. Current Opinion in Oncology 34, 243 (2022).

42. Lockhart, M. E. Separating the Benign from the Deadly: Active Surveillance of Oncocytoma after Biopsy. Radiology (2023) doi:10.1148/radiol.223108.

43. Preoperatively Misclassified, Surgically Removed Benign Renal Masses: A Systematic Review of Surgical Series and United States Population Level Burden Estimate | Journal of Urology. https://www.auajournals.org/doi/abs/10.1016/j.juro.2014.07.102.

44. Díaz-Montero, C. M., Rini, B. I. & Finke, J. H. The immunology of renal cell carcinoma. Nat Rev Nephrol 16, 721–735 (2020).

45. Chen, X. et al. Identifying tumor antigens and immune subtypes of renal cell carcinoma for immunotherapy development. Frontiers in Immunology 13, 1037808 (2022).

46. Hakimi, A. A. et al. An Integrated Metabolic Atlas of Clear Cell Renal Cell Carcinoma. Cancer cell 29, 104 (2016).

47. Wang, Y. et al. G3BP1 promotes tumor progression and metastasis through IL-6/G3BP1/STAT3 signaling axis in renal cell carcinomas. Cell Death Dis 9, 1–13 (2018).

48. Liu, Q. et al. Frequent Epigenetic Suppression of Tumor Suppressor Gene Glutathione Peroxidase 3 by Promoter Hypermethylation and Its Clinical Implication in Clear Cell Renal Cell Carcinoma. International Journal of Molecular Sciences 16, 10636–10649 (2015).

49. Ning, X.-H., et al. Association between FBP1 and hypoxia-related gene expression in clear cell renal cell carcinoma. Oncol Lett 11, 4095–4098 (2016).

50. Dondeti, V. R. et al. Integrative Genomic Analyses of Sporadic Clear Cell Renal Cell Carcinoma Define Disease Subtypes and Potential New Therapeutic Targets. Cancer Research 72, 112–121 (2012).

51. Li, B. et al. Fructose-1,6-bisphosphatase opposes renal carcinoma progression. Nature 513, 251–255 (2014).

52. Liu, T. et al. Hypoxia-induced PLOD2 promotes clear cell renal cell carcinoma progression via modulating EGFR-dependent AKT pathway activation. Cell Death Dis 14, 1–15 (2023).

53. Lidgren, A., Bergh, A., Grankvist, K., Rasmuson, T. & Ljungberg, B. Glucose transporter-1 expression in renal cell carcinoma and its correlation with hypoxia inducible factor-1α. BJU International 101, 480–484 (2008).

54. Betsunoh, H. et al. Clinical Significance of 18F-fluorodeoxyglucose and Glucose Transporter 1 mRNA in Clear Cell Renal Cell Carcinoma. Anticancer Research 41, 5179–5188 (2021).

55. Gasparre, G. et al. Disruptive mitochondrial DNA mutations in complex I subunits are markers of oncocytic phenotype in thyroid tumors. Proceedings of the National Academy of Sciences 104, 9001–9006 (2007).

56. Meierhofer, D. et al. Mitochondrial DNA mutations in renal cell carcinomas revealed no general impact on energy metabolism. Br J Cancer 94, 268–274 (2006).

57. Wang, Y. et al. Multi-omic profiling of intraductal papillary neoplasms of the pancreas reveals distinct expression patterns and potential markers of progression. 2024.07.07.602385 Preprint at 10.1101/2024.07.07.602385 (2024).

58. Zhou, Y. et al. Metascape provides a biologist-oriented resource for the analysis of systems-level datasets. Nat Commun 10, 1523 (2019).

59. Ashburner, M. et al. Gene Ontology: tool for the unification of biology. Nat Genet 25, 25–29 (2000).

59. The Gene Ontology Consortium et al. The Gene Ontology knowledgebase in 2023. Genetics 224, iyad031 (2023).

